# A rapidly deployable CRISPR–Cas3 diagnostic platform for emerging RNA viruses

**DOI:** 10.64898/2026.08.25.746999

**Authors:** Jun Nakamura, Kaya Miyazaki-Iida, Shotaro Torii, Masaaki Kitajima, Kouya Mikamo, Tetsutaro Kimihira, Lise Morimoto, Hamza Ashayqa, Jumpei Ito, Kohei Takeshita, Shingo Kosugi, Yasutaka Minegishi, Mutsumi Ito, Rika Hirano, Saeko Ishida, Kazuto Yoshimi, Peter J. Halfmann, Yoshihiro Kawaoka, Tomoji Mashimo

**Affiliations:** Division of Animal Genetics, Laboratory Animal Research Center, The Institute of Medical Science, The University of Tokyo, Tokyo 108-8639, Japan; Department of Urban Engineering, School of Engineering, The University of Tokyo, Tokyo 113-8656, Japan; Laboratory of International Wastewater-based Epidemiology, Research Center for Water Environment Technology, School of Engineering, The University of Tokyo, Tokyo 113-0032, Japan; Laboratory of Virus Informatics, Department of Biological Informatics, Bioinformatics Center, Research Institute for Microbial Diseases, The University of Osaka, Osaka 565-0871, Japan; Life Science Research Infrastructure Group, Advanced Photon Technology Division, RIKEN SPring-8 Center, Hyogo 679-5148, Japan; Research Reagents Group, Research and Development Section 3, Research and Development Department, NIPPON GENE CO., LTD., Toyama 930-0834, Japan; Division of Systems Virology, Department of Microbiology and Immunology, The Institute of Medical Science, The University of Tokyo, Tokyo 108-8639, Japan; Laboratory Animal Science, Graduate School of Medicine and Faculty of Medicine, Kyoto University, Kyoto, Japan; Department of Pathobiological Sciences, School of Veterinary Medicine, University of Wisconsin–Madison, Madison, WI 53706, USA; Division of Virology, Department of Microbiology and Immunology, The Institute of Medical Science, The University of Tokyo, Tokyo 108-8639, Japan; The University of Tokyo Pandemic Preparedness, Infection and Advanced Research Center (UTOPIA), Tokyo, Japan; Division of Genome Engineering, Center for Experimental Medicine and Systems Biology, The Institute of Medical Science, The University of Tokyo, Tokyo 108-8639, Japan

## Abstract

Rapidly converting viral genome information into deployable molecular tests remains a major challenge in outbreak preparedness. We developed CONAN-SWIFT (Simple Workflow for Isothermal Field Testing), a sequence-to-test platform that integrates computational assay design, reverse-transcription loop-mediated isothermal amplification, CRISPR–Cas3 detection, reagent lyophilization and lateral-flow readout. Sequence-guided assays for Andes virus and Bundibugyo virus were established within approximately three weeks and extended to four additional filoviruses. A web-based designer supported crRNA selection, and systematic RT-LAMP primer optimization improved amplification performance. Recombinant Escherichia coli-expressed Cascade enabled standardized preparation of lyophilized Cas3-detection reagents, which were combined with a battery-operated isothermal device. The portable system detected as few as 10 input RNA copies per reaction within approximately 40 min. It also detected viral RNA and biologically contained, replication-incompetent Ebola virus in spiked human blood and concentrated wastewater. These findings establish the analytical feasibility of a rapidly adaptable CRISPR–Cas3 engineering framework for decentralized detection of emerging RNA viruses.

## Introduction

In May 2026, an outbreak of Andes virus (ANDV)-associated hantavirus pulmonary syndrome linked to travel aboard the expedition cruise ship M/V Hondius resulted in cases and contacts across multiple countries, requiring medical evacuation, repatriation, coordinated international contact tracing, isolation and quarantine [1,2]. During the same period, an outbreak of Ebola disease caused by Bundibugyo virus (BDBV) affected the Democratic Republic of the Congo and Uganda and was declared a Public Health Emergency of International Concern [3–5]. A report describing the 2026 Ugandan index case further highlighted how nonspecific early symptoms, cross-border movement and delayed recognition can impede timely diagnosis and outbreak control [5]. Together, these events highlighted a critical vulnerability in outbreak preparedness: suspected cases may present at hospitals, isolation facilities or international points of entry before pathogen-specific diagnostic tests become locally available.

Reverse-transcription quantitative PCR (RT-qPCR) remains the reference laboratory method for detecting ebolaviruses and hantaviruses but generally requires specialized equipment, trained personnel, reliable electricity and controlled sample transport [6,7]. Rapid antigen tests are more readily deployable but may show reduced sensitivity at low viral loads and often require confirmatory molecular testing [7,8]. Moreover, the development, validation and manufacture of pathogen-specific diagnostic tests can take weeks to months, creating a critical diagnostic gap during the early phase of an outbreak [9,10]. Isothermal amplification and CRISPR-based diagnostics (CRISPR-Dx) provide complementary approaches because they enable sensitive and sequence-specific nucleic acid detection and are compatible with portable fluorescence or lateral-flow readouts [11–15]. We previously developed Cas3-operated nucleic acid detection (CONAN), which exploits target-dependent collateral DNA cleavage by the type I-E CRISPR–Cas3 system for rapid viral detection [16], and subsequently developed Kairo-CONAN, a portable testing system combining lyophilized reagents with a simplified heating device [17].

Nevertheless, translating newly available viral genome sequences into deployable molecular tests requires coordinated solutions for target selection, amplification design, reagent production, stabilization, incubation and readout. Here we developed CONAN-SWIFT, an integrated sequence-to-test engineering framework combining computational crRNA selection, RT-LAMP amplification, Cas3-operated nucleic acid detection, lyophilized reagents and lateral-flow readout. We used contemporaneous ANDV and BDBV outbreaks as test cases, then examined whether the same workflow could be extended across a panel of medically important filoviruses and operated in blood and wastewater matrices. By linking assay design to a portable testing format, CONAN-SWIFT is intended to shorten the path from pathogen sequence availability to an analytically validated prototype, consistent with the objectives of the 100 Days Mission [18].

## Results

### Sequence-guided development of ANDV- and BDBV-specific RT-LAMP–CONAN assays

Following the availability of outbreak-associated ANDV genome sequences and the publication of interim diagnostic guidance on 15 May 2026 [1,2,19], comparison of ANDV with related pathogenic hantaviruses identified conserved regions suitable for ANDV-specific detection within the nucleoprotein-coding region of the small (S) genomic segment (Fig. 1a and Extended Data Fig. 1a). Three candidate crRNAs were evaluated against synthetic double-stranded DNA targets using fluorescence-based CONAN [16,20], with crRNA2 and crRNA3 showing the strongest target-dependent activity (Fig. 1b and Extended Data Fig. 1b). The synthetic in vitro-transcribed ANDV RNA used for assay development was quality-controlled by TapeStation analysis (Fig. 1c). RT-LAMP primer sets encompassing the crRNA2 and crRNA3 target regions were then evaluated through three successive screening rounds (Fig. 1d and Extended Data Fig. 1c,d). The selected RT-LAMP products were detected by CONAN using a lateral-flow assay (LFA), with no detectable test lines in the corresponding no-template controls (Fig. 1e).crRNA2 and crRNA3 showed the highest activity at 10 and 100 nM, respectively, and were therefore carried forward for LFA-based evaluation. The crRNA2-based assay (ANDV CL#2-3), which produced the stronger target-dependent LFA signal, was selected for subsequent experiments (Fig. 1b,e). Thus, a prototype ANDV-specific RT-LAMP–CONAN assay was established within 25 days.

**Figure 1.**
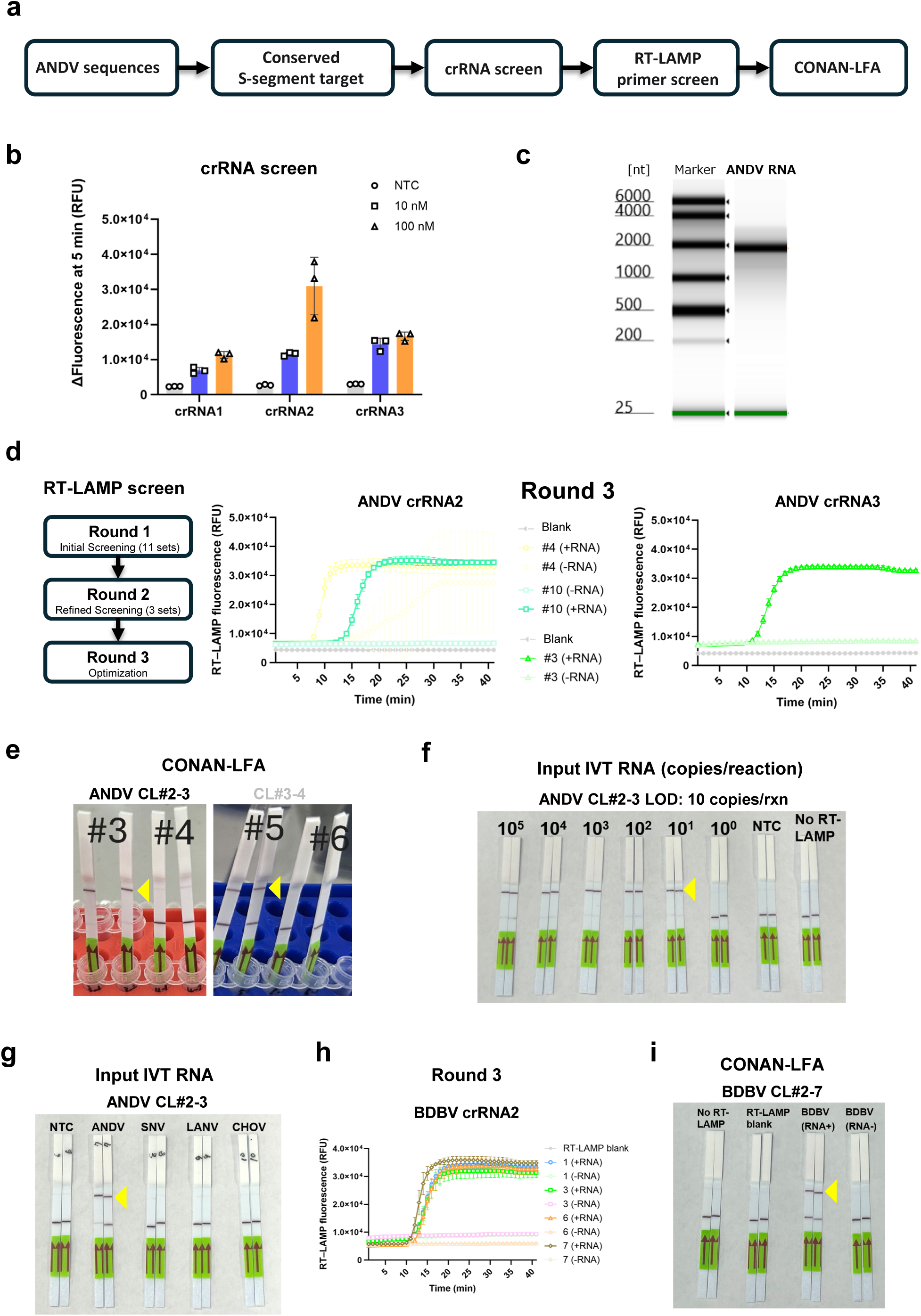
Rapid sequence-guided development of ANDV- and BDBV-specific RT-LAMP–CONAN assays. a, Sequence-guided workflow for development of the ANDV RT-LAMP–CONAN assay. b, Fluorescence-based screening of candidate ANDV crRNAs using synthetic double-stranded DNA targets. c, Representative TapeStation analysis of purified in vitro-transcribed (IVT) ANDV RNA used for assay development. d, Three-round screening of RT-LAMP primer sets targeting the ANDV crRNA2 and crRNA3 regions, resulting in selection of ANDV CL#2-3 and CL#3-4. e, Lateral-flow detection of synthetic IVT ANDV RNA using the selected assays. f, Analytical sensitivity of ANDV CL#2-3 using serially diluted IVT ANDV RNA, showing detection down to 10 copies per reaction. g, Analytical specificity of ANDV CL#2-3 against IVT RNAs from ANDV and the related hantaviruses Sin Nombre virus (SNV), Laguna Negra virus (LANV) and Choclo virus (CHOV), assessed by lateral-flow assay. h, Final-round RT-LAMP screening for the BDBV crRNA2 target region. i, Lateral-flow detection of synthetic IVT BDBV RNA using the selected BDBV CL#2-7 assay. NTC, no-template control; IVT, in vitro transcription; LFA, lateral-flow assay. Yellow arrowheads indicate positive test (T) lines.

We next evaluated the analytical sensitivity and specificity of the selected ANDV assay by LFA. Using serially diluted synthetic in vitro-transcribed ANDV RNA, ANDV CL#2-3 detected 10 RNA copies per reaction under the tested conditions (Fig. 1f). Because ANDV is the only hantavirus for which person-to-person transmission has been documented [21,22], discrimination from related pathogenic hantaviruses is particularly relevant to infection-control measures and contact management. No cross-reactivity was observed with synthetic in vitro-transcribed RNAs derived from Sin Nombre virus (SNV), Laguna Negra virus (LANV) or Choclo virus (CHOV) (Fig. 1g).

We next applied the same workflow to Bundibugyo virus (BDBV), which caused an Ebola disease outbreak affecting the Democratic Republic of the Congo and Uganda during the same period [3,4]. Comparative sequence analysis identified BDBV-conserved regions that were divergent from other medically important filoviruses (Extended Data Fig. 2a,b). The synthetic in vitro-transcribed BDBV RNA used for assay development was assessed by TapeStation analysis (Extended Data Fig. 2c). Candidate crRNAs were evaluated using fluorescence-based CONAN, and crRNA2 was selected for assay development (Extended Data Fig. 2d). RT-LAMP primer sets encompassing the crRNA2 target region were screened using synthetic in vitro-transcribed BDBV RNA, with the final screening round shown in Fig. 1h. The selected crRNA–primer combination, BDBV CL#2-7, generated a target-dependent test line by LFA, whereas the no-template control remained negative (Fig. 1i). A prototype BDBV-specific RT-LAMP–CONAN assay was established within 21 days. Together, these findings demonstrate that the sequence-guided workflow can be adapted to distinct emerging RNA viruses within weeks.

### Computational and experimental optimization of RT-LAMP–CONAN assays

Although the initial BDBV-specific RT-LAMP–CONAN assay (BDBV CL#2-7) enabled target detection, its sensitivity was lower than that of the ANDV CL#2-3 assay, indicating a need for further optimization. We therefore improved computational crRNA selection and RT-LAMP primer design. First, we developed CONAN-SWIFT Designer, a web-based application that facilitates crRNA selection from viral genome sequences (Fig. 2a and Supplementary Protocol 1). The application identifies conserved target-specific regions and ranks candidate crRNAs according to target conservation, non-target specificity and compatibility with RT-LAMP primer design. Potential off-target matches can also be assessed using sequence-similarity searches, facilitating candidate selection for experimental validation (Extended Data Fig. 3).

**Figure 2.**
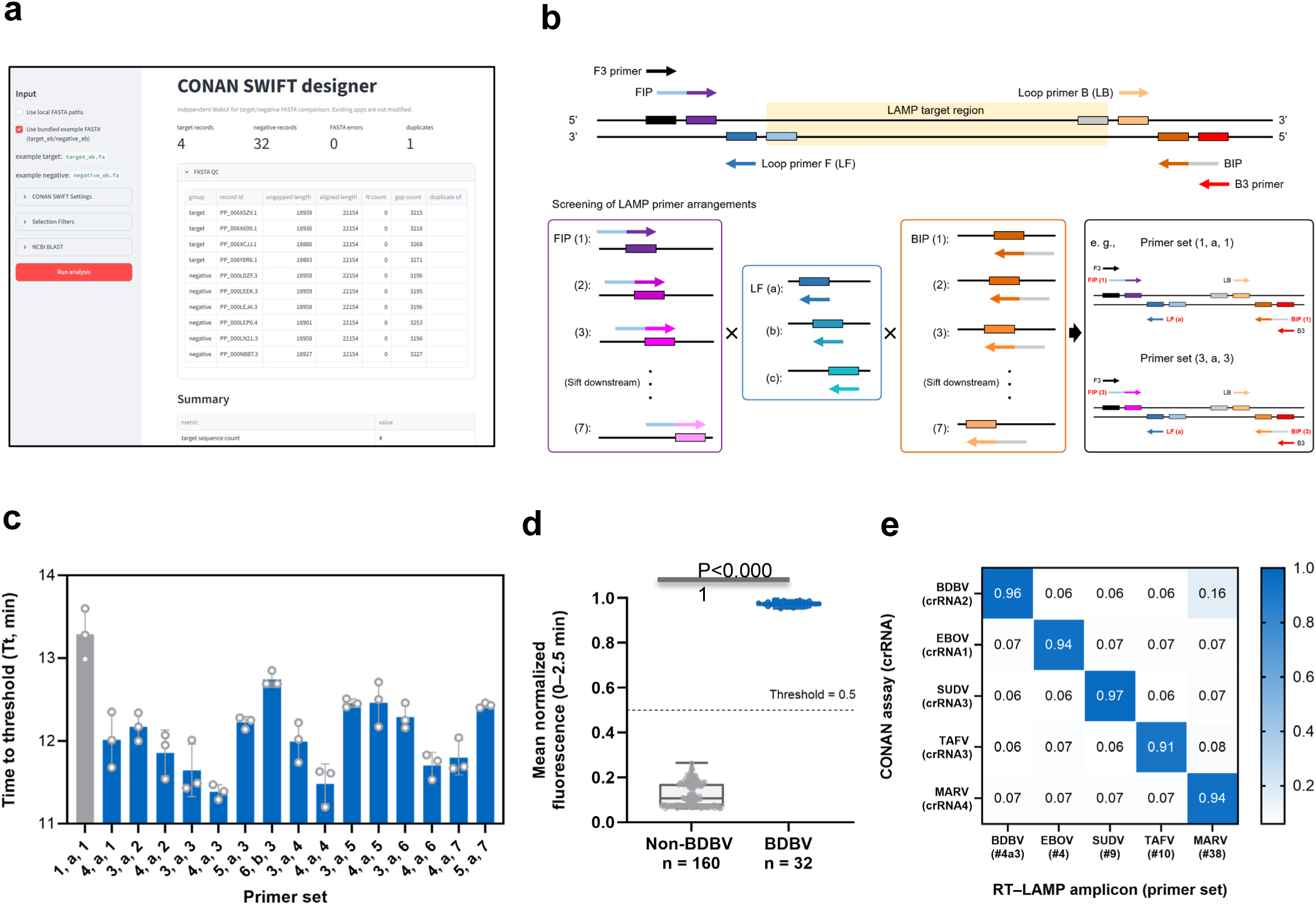
Computational and experimental optimization of RT-LAMP–CONAN assays. a, Schematic of the CONAN-SWIFT Designer workflow for sequence-guided selection of target-specific crRNAs and compatible RT-LAMP target regions from viral genome sequences. Candidate crRNAs are prioritized according to target conservation, non-target specificity, off-target risk and compatibility with RT-LAMP primer design. b, Systematic optimization of RT-LAMP primer architecture for the BDBV CL#2-7 assay by repositioning the F2- and B2-binding sites and corresponding loop-primer positions to generate primer sets with different F2–B2 spans. c, Comparison of RT-LAMP amplification performance among BDBV primer configurations. Screening of 147 configurations identified primer set 4a3, which reduced the threshold time (Tt) by 1.91 min (14.3%) relative to the original 1a1 primer set and was selected as BDBV CL#2-7-4a3. d, Analytical specificity of the optimized BDBV CL#2-7-4a3 assay, assessed by normalized fluorescence at 2.5 min using synthetic IVT RNAs from BDBV (n = 32) and non-BDBV filoviruses (EBOV, SUDV, TAFV and MARV; n = 160). The dashed line indicates the positivity threshold (0.5). Two-sided exact Mann–Whitney U-test, P<0.0001. e, Cross-reactivity matrix of virus-specific RT-LAMP–CONAN assays for BDBV, EBOV, SUDV, TAFV and MARV. Each assay detected its corresponding synthetic IVT RNA without detectable cross-reactivity with the other four filovirus targets under the tested conditions. NTC, no-template control.

We next optimized RT-LAMP primer design for BDBV RNA amplification by generating multiple candidate FIP, BIP and loop-forward (LF) primers within the aligned target region and systematically testing 147 primer combinations (Fig. 2b). Primer set 4a3 reduced the threshold time (Tt) by 1.91 min (14.3%) relative to the original 1a1 set and showed lower inter-replicate variability (Fig. 2c). BDBV CL#2-7-4a3 was therefore selected for subsequent experiments. The optimized BDBV assay showed no cross-reactivity with synthetic in vitro-transcribed RNAs derived from four other filoviruses—Ebola virus (EBOV), Sudan virus (SUDV), Taï Forest virus (TAFV) and Marburg virus (MARV)—using LFA readout (Fig. 2d). We then used CONAN-SWIFT Designer to design virus-specific crRNAs and RT-LAMP primers for these four viruses and selected lead crRNA–primer combinations by fluorescence-based crRNA screening and RT-LAMP primer screening (Extended Data Fig. 4a,b). When the resulting five RT-LAMP–CONAN assays, including the BDBV assay, were evaluated as a panel, each assay detected only its corresponding viral RNA, with no cross-reactivity against the other four viruses under the tested conditions (Fig. 2e). These results demonstrate that the sequence-guided design strategy facilitates the development of specific RT-LAMP–CONAN assays capable of distinguishing medically important filoviruses (Supplementary Protocol 2).

### Integration of lyophilized reagents into the portable CONAN-SWIFT system

To translate the optimized assays into a standardized and portable format, we used target-specific crRNA-loaded Cascade ribonucleoprotein complexes expressed and purified from Escherichia coli (ecCascade) [16,20], replacing the in vitro-expressed Cascade (ivCascade) used during initial assay development. Cas3 and crRNA-loaded ecCascade were formulated as ready-to-use lyophilized CONAN reagents, as previously described [17]. Using EGFP as a model target, the reconstituted lyophilized formulation retained target-dependent CONAN activity comparable to that of the corresponding liquid formulation (Fig. 3a). The lyophilized Cas3–Cascade reagents also retained target-dependent collateral-cleavage activity after storage at 25 °C for up to 70 days (Extended Data Fig. 5).

**Figure 3.**
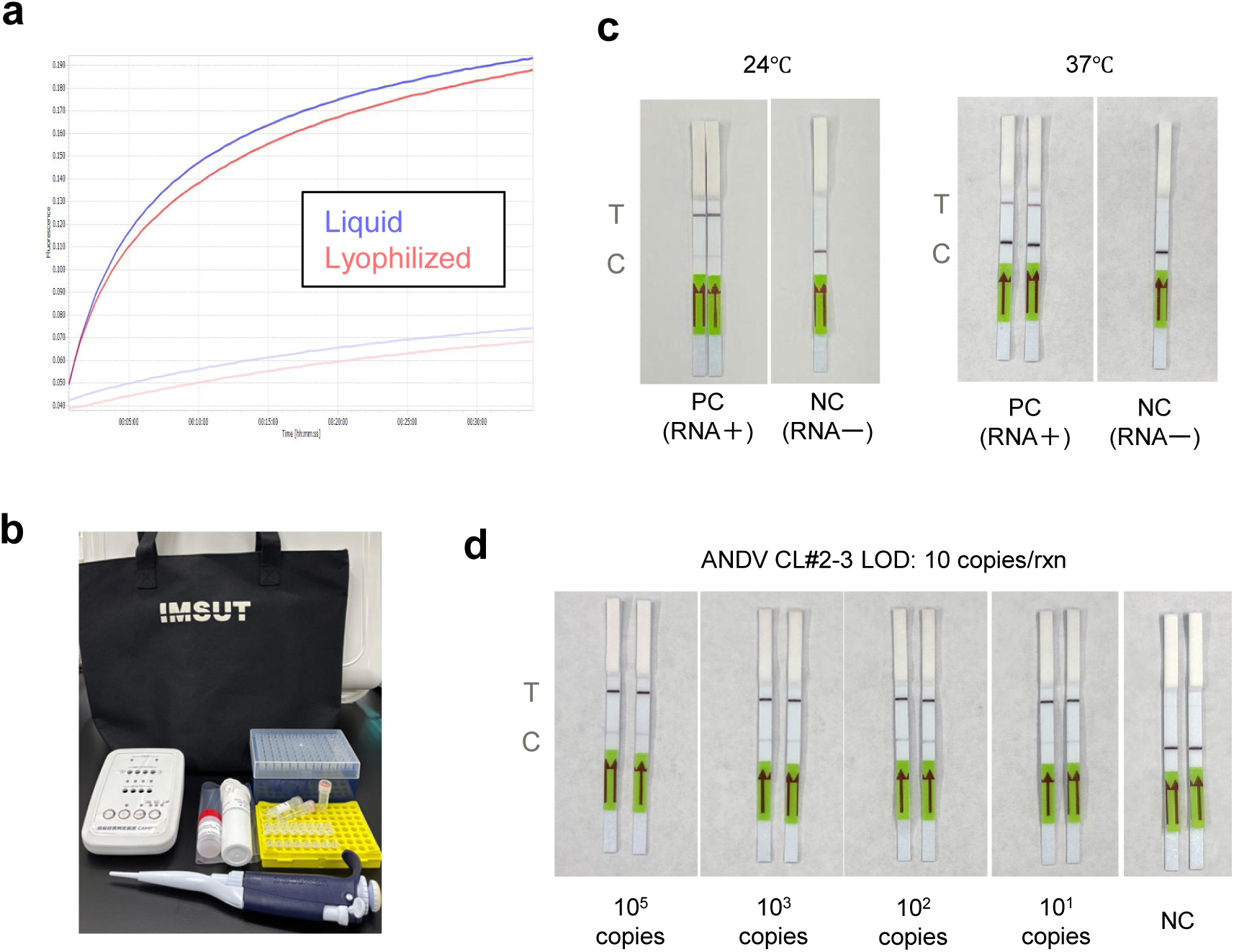
Development of the portable CONAN-SWIFT system for on-site viral RNA detection. a, Comparison of liquid and lyophilized CONAN reagents using EGFP as a model target. Reconstituted lyophilized Cas3 and crRNA-loaded ecCascade retained target-dependent CONAN activity. Red, lyophilized reagents; blue, liquid reagents; dark colours, target-containing reactions; pale colours, reagent-only controls without template. b, Configuration of the portable CONAN-SWIFT system, integrating lyophilized RT-LAMP and CONAN reagents, lateral-flow readout and the battery-operated CAMP isothermal device. The complete workflow comprises 30 min of RT-LAMP amplification, 5 min of CONAN detection and 3 min of lateral-flow development, enabling visual detection within 40 min (Supplementary Protocol 3 and Video 1). c, Detection of ANDV RNA using CONAN-SWIFT at ambient temperatures of 24 °C and 37 °C. d, Analytical sensitivity of CONAN-SWIFT using serially diluted synthetic in vitro-transcribed ANDV RNA, showing detection down to 10 RNA copies per reaction. C, control line; T, test line; NC, negative control; PC, positive control.

We next combined the lyophilized CONAN reagents with the commercially available lyophilized HeatAct LAMP MASTER kit (NIPPON GENE) for RT-LAMP amplification. Commercial lateral-flow strips were compared for visual discrimination between positive and negative CONAN reactions, and the strip providing the clearest readout was selected for subsequent experiments (Extended Data Fig. 6). Portable isothermal amplification devices were also compared, and the compact, lightweight, battery-operated CAMP device (NIPPON GENE) was selected for field-compatible incubation. These components were integrated into CONAN-SWIFT (Simple Workflow for Isothermal Field Testing) (Fig. 3b).

Using the ANDV CL#2-3 assay, CONAN-SWIFT produced clear target-dependent lateral-flow signals at both 24 °C and 37 °C, demonstrating robust operation across the tested ambient temperatures (Fig. 3c). Serial dilution of synthetic in vitro-transcribed ANDV RNA showed detection down to 10 RNA copies per reaction (Fig. 3d). The complete portable workflow comprises reagent reconstitution, 30 min of RT-LAMP amplification, 5 min of CONAN detection and 3 min of lateral-flow development, enabling visual detection within 40 min. The workflow is detailed in Supplementary Protocol 3 and demonstrated in Supplementary Video 1.

### Extraction-free detection of viral targets in human whole blood

Rapid detection of viral targets in blood is important for point-of-care testing of viral haemorrhagic fevers [5]. We therefore established an extraction-free CONAN-SWIFT workflow for human whole blood (Fig. 4a). A crRNA targeting human RPP20 was first validated by fluorescence-based CONAN using RT-LAMP products (Fig. 4b). To assess matrix-associated inhibition, synthetic RPP20 RNA was analysed in undiluted whole blood and after dilution with water. Among the tested dilutions, a 1:3 blood-to-water ratio (fourfold final dilution) provided the best balance between alleviating blood-associated inhibition and retaining sufficient target concentration and analytical sensitivity, and was therefore selected for subsequent analyses. (Fig. 4c).

**Figure 4.**
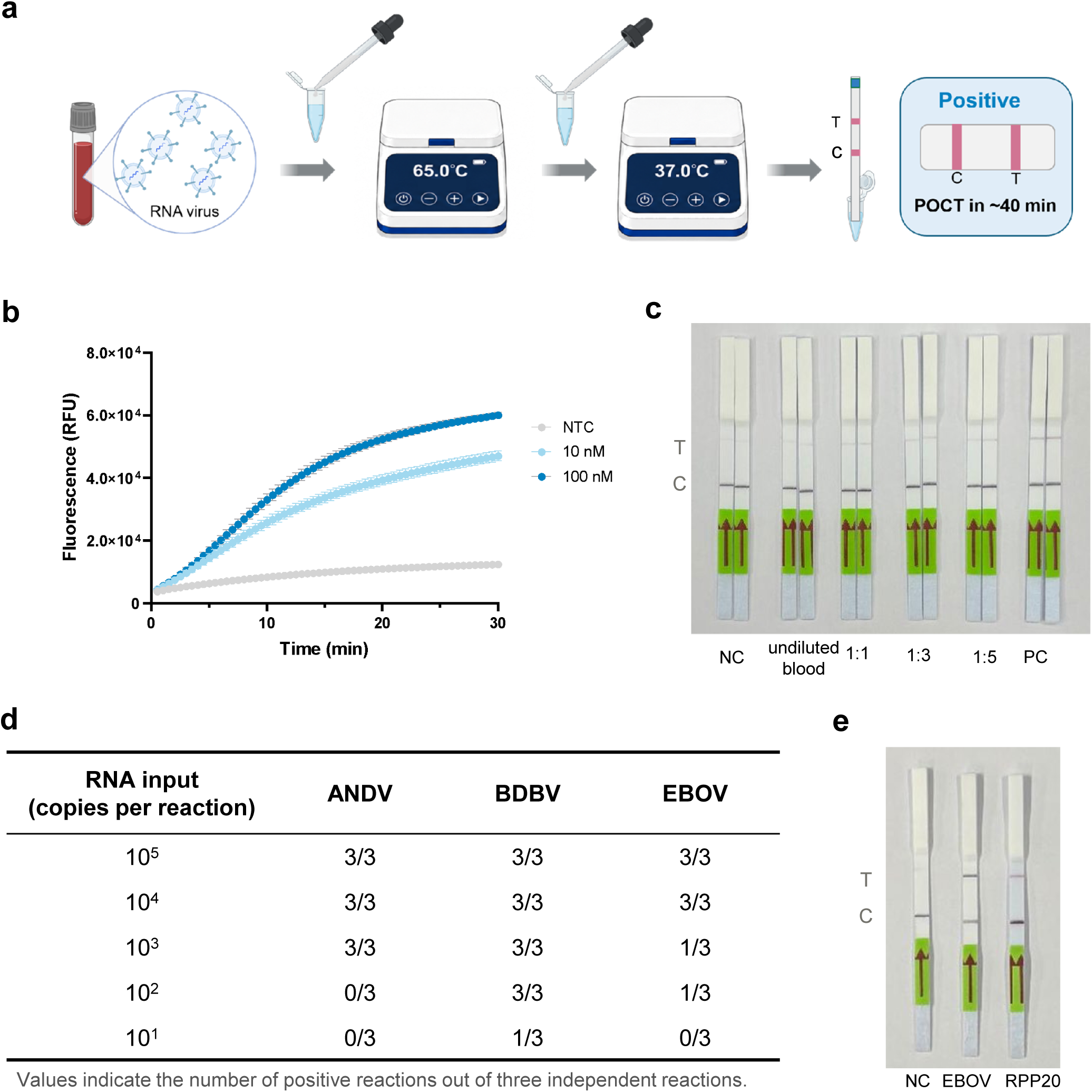
Detection of viral targets in human whole blood using CONAN-SWIFT. a, Schematic overview of extraction-free point-of-care testing of human whole blood using CONAN-SWIFT, comprising sample dilution, RT-LAMP amplification, CONAN detection and visual lateral-flow readout. b, Validation of a crRNA targeting human RPP20 by fluorescence-based CONAN using RT-LAMP products. c, Effect of whole-blood dilution on RPP20 detection. Synthetic RPP20 RNA (10⁵ copies per reaction) was spiked into whole blood, which was tested undiluted or after dilution with dH₂O at the indicated ratios, followed by lateral-flow readout. d, Detection of serially diluted synthetic ANDV, BDBV and EBOV RNAs in whole blood. Values indicate positive reactions among three independent reactions and represent detection frequency rather than a formally determined limit of detection; representative lateral-flow strips are shown in Extended Data Fig. 7a. e, Detection of biologically contained EBOV ΔVP30 and endogenous RPP20 in whole blood from a healthy donor after 1:3 dilution with dH₂O. RPP20 served as an internal sample control. T, test line; C, control line; NC, negative control; PC, positive control; NTC, no-template control.

We next evaluated viral target detection using the same extraction-free sample preparation. Serially diluted synthetic in vitro-transcribed ANDV, BDBV and EBOV RNAs were analysed in whole blood, with detection frequency assessed across three independent reactions at each input level (Fig. 4d and Extended Data Fig. 7a). We further evaluated biologically contained, replication-incompetent Ebola virus lacking the VP30 gene (EBOV ΔVP30) [23]. EBOV ΔVP30 was analysed in whole blood after 1:3 dilution with water, with endogenous RPP20 serving as an internal sample control. A positive EBOV ΔVP30 signal was obtained in diluted whole blood (Fig. 4e). These findings support the analytical feasibility of extraction-free CONAN-SWIFT testing in whole blood (Supplementary Protocol 4), although validation using clinical specimens will be required.

### Detection of emerging viral targets in wastewater

Wastewater surveillance enables community-level monitoring and early detection of emerging viruses [24,25]. We therefore evaluated CONAN-SWIFT in airport and urban wastewater using a workflow comprising sample concentration, nucleic acid extraction, RT-LAMP amplification, CONAN detection and lateral-flow readout (Fig. 5a). As an endogenous wastewater-associated control, we targeted pepper mild mottle virus (PMMoV), a widely used indicator of human faecal contamination [26]. Endogenous PMMoV RNA was detected in wastewater samples by CONAN-SWIFT (Fig. 5b).

**Figure 5.**
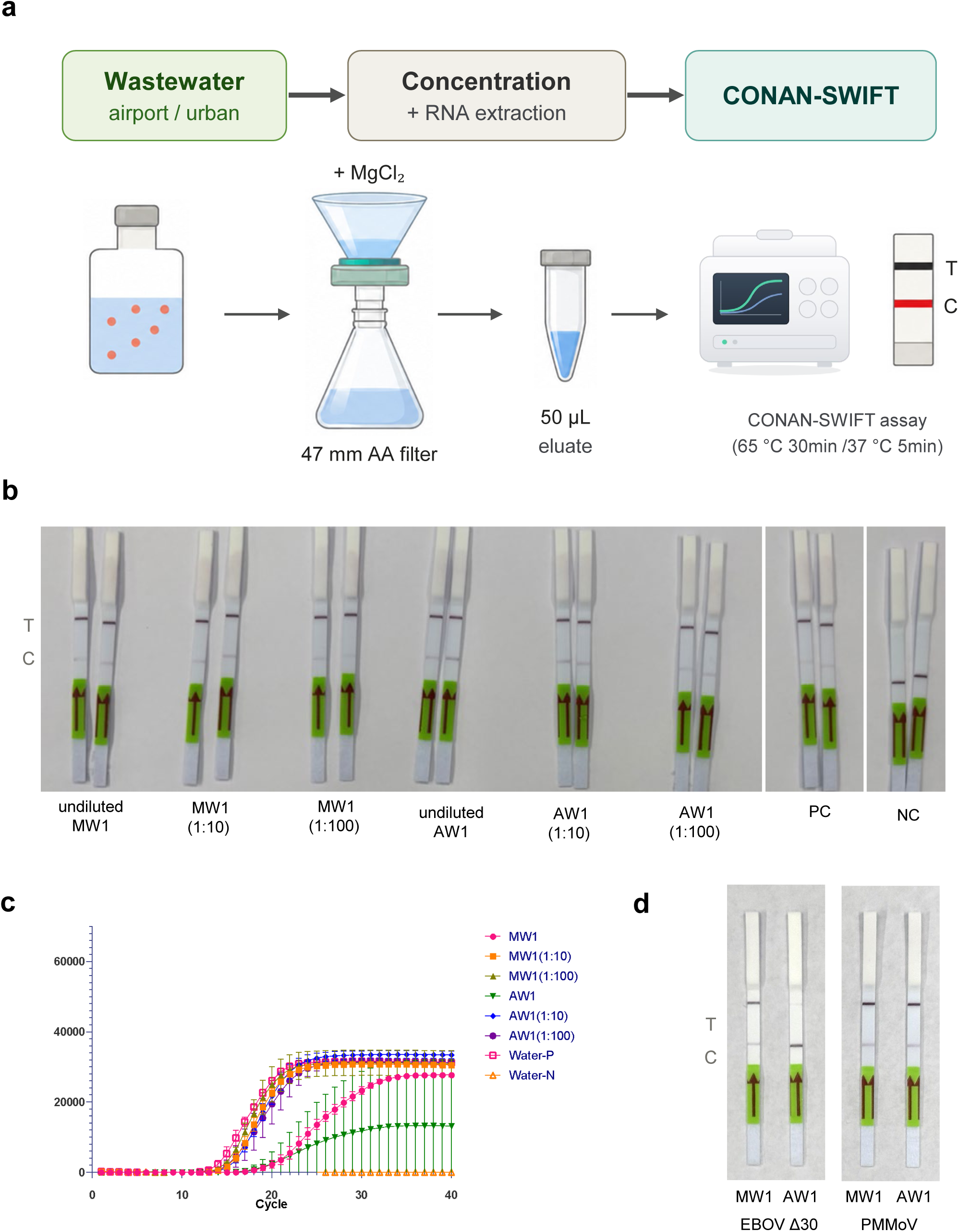
Application of CONAN-SWIFT to viral target detection in wastewater. a, Schematic overview of the CONAN-SWIFT workflow for wastewater analysis, comprising sample concentration, nucleic acid extraction, RT-LAMP amplification, CONAN detection and lateral-flow readout. b, Detection of endogenous pepper mild mottle virus (PMMoV) RNA in airport and urban wastewater samples using CONAN-SWIFT. c, Effect of dilution on RT-LAMP amplification of wastewater-derived RNA. Undiluted, 1:10-diluted and 1:100-diluted RNA extracts from representative municipal wastewater (MW1) and airport wastewater (AW1) samples were analysed by real-time RT-LAMP. d, Detection of biologically contained EBOV ΔVP30 spiked into unconcentrated airport and urban wastewater before sample concentration and nucleic acid extraction. Endogenous PMMoV was detected in parallel as a wastewater-associated process control. C, control line; T, test line; NC, negative control; PC, positive control.

We next examined matrix-associated inhibition using wastewater-derived nucleic acid extracts. Dilution of representative municipal wastewater (MW1) and airport wastewater (AW1) extracts improved RT-LAMP amplification relative to undiluted material (Fig. 5c). Finally, biologically contained, replication-incompetent EBOV ΔVP30 [23] was spiked into airport and urban wastewater samples following sample concentration and nucleic acid extraction. Under the tested conditions, EBOV ΔVP30 was detected in the urban wastewater sample by CONAN-SWIFT, with endogenous PMMoV detected in parallel as a wastewater-associated process control (Fig. 5d). Analytical sensitivity of synthetic BDBV and EBOV RNAs in wastewater-derived extracts was evaluated separately (Extended Data Fig. 7b). Together, these results support the analytical feasibility of CONAN-SWIFT for viral target detection in complex wastewater matrices (Supplementary Protocol 5).

## Discussion

A central engineering challenge in outbreak diagnostics is not only achieving analytical sensitivity, but also rapidly converting pathogen sequence information into a manufacturable and portable test. Single-effector Cas12- and Cas13-based diagnostics have demonstrated sensitive, field-compatible nucleic acid detection [12–15,28–31]. CONAN-SWIFT provides a complementary sequence-to-test framework based on the multisubunit type I-E CRISPR–Cas3 system and its target-dependent collateral single-stranded DNA cleavage activity [16,32]. The use of a common computational and experimental workflow across ANDV, BDBV and four additional filoviruses indicates that the platform can be reconfigured for phylogenetically distinct emerging RNA viruses within weeks.

The integration of standardized recombinant Cascade, lyophilized reagents, battery-operated isothermal incubation and lateral-flow readout addresses several requirements for testing outside centralized laboratories. Previous portable RT-LAMP and CRISPR systems have established the feasibility of minimally instrumented molecular testing [27–30]; the distinguishing feature of CONAN-SWIFT is the connection between sequence-based assay design and a portable Cas3 testing format. Detection in spiked whole blood and wastewater further demonstrates tolerance of clinically and environmentally relevant matrices. However, the study used synthetic RNA, spiked matrices and biologically contained EBOV ΔVP30 rather than specimens from infected patients [23]. It therefore establishes analytical feasibility, not clinical sensitivity, specificity or predictive performance. Detection limits require confirmation with larger replicate numbers and formal statistical analyses, and direct comparisons with RT-qPCR, antigen tests and Cas12- or Cas13-based diagnostics are needed to define relative performance, cost and operational value.

The current two-step workflow requires transfer of amplified material from RT-LAMP to the CONAN reaction, increasing handling and the risk of amplicon contamination. Further development should prioritize a sealed, single-use architecture that preserves sequential amplification and Cas3 detection without opening the reaction vessel. Automated sample preparation, reagent reconstitution, fluid transfer, incubation and objective signal interpretation would reduce operator-dependent variability and exposure risk. One-pot and minimally instrumented CRISPR systems demonstrate that such integration is feasible [29,30], although the biochemical requirements of RT-LAMP and CONAN must be reconciled. Until a closed system is developed and clinically validated, specimens from patients with suspected viral haemorrhagic fever should be tested by trained healthcare or laboratory personnel under appropriate biosafety conditions.

In conclusion, CONAN-SWIFT links computational assay design, experimental screening, reagent stabilization and portable visual detection in a single CRISPR–Cas3 development framework. Its principal contribution is a systematic pathway for generating deployable prototypes rather than a single pathogen-specific diagnostic product. Future work should focus on closed-system automation, quality-controlled manufacturing, shelf-life and transport stability, and prospective evaluation with authentic clinical specimens. Multi-site field studies should assess usability, biosafety and environmental surveillance under outbreak-relevant conditions and determine whether CONAN-SWIFT can complement centralized RT-qPCR for earlier detection of emerging viral threats.

## Methods

### Sequence analysis and crRNA target selection

Reference nucleotide sequences for each viral target and, where applicable, related non-target viruses were obtained from the NCBI GenBank/RefSeq databases (accession and version numbers are provided in Supplementary Table 1). Multiple-sequence alignments were generated from the downloaded sequences and inspected manually. For ANDV and the filovirus assays, candidate crRNA target regions were selected by visual inspection of the alignments, prioritizing regions with limited sequence variation among target sequences across an approximately 200-nt local window while retaining sequence differences from related non-target viruses. Candidate Cascade target sites were restricted to 32-nt protospacers adjacent to a 5′-AAG-3′ PAM. Sequence variation within the PAM-proximal 10 nt of each protospacer was examined particularly closely, and sites showing minimal variation in this region were preferentially selected. Candidate regions containing ambiguous nucleotides or alignment gaps within sequences required for crRNA recognition were excluded.

For PMMoV, candidate crRNAs were identified using CONAN-SWIFT Designer as described below. For the human RNase P internal control (RPP20/POP7), a Cascade crRNA target was selected within the amplicon defined by the previously reported RT-LAMP primer set of Curtis et al. (2018), by identifying a 32-nt protospacer adjacent to a 5′-AAG-3′ PAM. The sequences of all crRNAs used in this study are provided in Supplementary Table 2.

### Computational design and ranking of Cas3-CONAN crRNAs

A custom Python-based web application, CONAN-SWIFT Designer, was developed to identify and rank candidate crRNAs from target and non-target FASTA sequences. Both DNA strands were searched for AAG PAMs, and adjacent 32-nucleotide sequences were extracted as candidate spacers. The eight PAM-proximal nucleotides were operationally defined as the seed region.

Candidates were evaluated based on conservation of the PAM, seed and spacer among target sequences; absence of PAM–seed or PAM–spacer matches in non-target sequences; sequence-similarity-based off-target risk; and the availability of conserved flanking regions suitable for RT-LAMP primer design. Candidates were excluded if the PAM was not conserved, the seed contained mismatches, the spacer contained ambiguous nucleotides or alignment gaps, or an exact PAM–seed match occurred in the non-target set. One or two PAM-distal mismatches were permitted with a scoring penalty.

Candidate crRNAs were ranked using a 100-point composite score incorporating target conservation, non-target specificity, off-target risk, sequence quality and suitability of approximately 200-nucleotide flanking regions for RT-LAMP primer design. The application was implemented in Python 3.12 using Streamlit and was used to prioritize candidates for experimental evaluation rather than to quantitatively predict CONAN activity.

### Plasmid construction

DNA template plasmids for in vitro transcription (IVT) were constructed using an in-house-modified derivative of the Template Vector (BspQ I) for T7 mRNA Synthesis (Takara Bio, cat. no. 6146), hereafter referred to as the pIVT backbone. To generate the pIVT backbone, a synthetic double-stranded DNA cassette (5′-GGGTCTTCGCTAGCGAAGACCT-3′), containing two oppositely oriented BbsI recognition sites separated by a 6-bp spacer, was ligated into the linearized cloning site between the 5′ and 3′ untranslated regions of the commercially supplied vector, thereby circularizing the vector. For target-fragment insertion, the pIVT backbone was linearized by overnight digestion with BbsI-HF (New England Biolabs, cat. no. R3539S) at 37 °C. Synthetic DNA fragments corresponding to ANDV-N, RPP20, BDBV-U, BDBV-D, EBOV-U, EBOV-D, SUDV-U, SUDV-D, TAFV-U, TAFV-D, MARV-U and MARV-D were obtained from Eurofins Genomics (Supplementary Table 1) and inserted into the linearized pIVT backbone using the In-Fusion Snap Assembly Master Mix (Takara Bio, cat. no. 638947). The assembled plasmids were transformed into NEB Stable Competent Escherichia coli (New England Biolabs, cat. no. C3040I) and purified using the PureLink HiPure Plasmid Filter Midiprep Kit (Invitrogen, cat. no. K210015). The resulting constructs were designated pIVT-ANDV-N, pIVT-RPP20, pIVT-BDBV-U, pIVT-BDBV-D, pIVT-EBOV-U, pIVT-EBOV-D, pIVT-SUDV-U, pIVT-SUDV-D, pIVT-TAFV-U, pIVT-TAFV-D, pIVT-MARV-U and pIVT-MARV-D. All plasmid inserts were verified by Sanger sequencing.

For the preparation of *in vitro*-expressed Cascade complexes, target-specific crRNA expression plasmids were constructed using the pACYC-Duet-1-derived vector pACYCDuet-1_crRNA-empty-BbsI, as described previously [16,20]. For each crRNA, complementary DNA oligonucleotides encoding the 32-nucleotide spacer and the appropriate cloning overhangs were synthesized by Eurofins Genomics (Supplementary Table 3). The complementary oligonucleotides were mixed at a final concentration of 100 μM each, heated at 95 °C for 5 min and gradually cooled to 25 °C at 0.1 °C s⁻¹. The annealed oligonucleotides were ligated into the BbsI-HF-digested pACYCDuet-1_crRNA-empty-BbsI vector using the DNA Ligation Kit (Takara Bio, cat. no. 6023). The resulting plasmids were designated pACYCDuet-1-[target abbreviation]-crRNA[number]. All constructs were verified by Sanger sequencing (Supplementary Table 4).

### In vitro transcription and purification of target RNAs

Linear DNA templates for IVT were prepared from the sequence-verified pIVT target plasmids either by restriction enzyme digestion or by PCR amplification using target-specific primers and Tks Gflex DNA Polymerase (Takara Bio, cat. no. R060B). PCR products were separated by agarose-gel electrophoresis and purified using the NucleoSpin Gel and PCR Clean-up kit (Macherey-Nagel, 740609.25). The sequences of the purified DNA templates were confirmed by Sanger sequencing.

Target RNAs were generated using the MEGAshortscript T7 Transcription Kit (Invitrogen, AM1354) according to the manufacturer’s instructions. Following transcription, residual template DNA was digested using the TURBO DNase supplied with the kit. RNA products were purified using the Monarch Spin RNA Cleanup Kit (50 µg; New England Biolabs, T2040L). RNA concentrations were measured using a NanoPhotometer N60 microvolume UV–visible spectrophotometer (Implen). The quality of IVT RNA preparations was qualitatively assessed using a 4150 TapeStation System and High Sensitivity RNA ScreenTape (Agilent Technologies, 5067-5579).

### Preparation of crRNA-loaded Cascade complexes

Target-specific crRNA-loaded Cascade complexes were prepared using sequence-verified crRNA expression plasmids (Supplementary Table 4), essentially as described previously [16,20]. Briefly, recombinant Cascade ribonucleoprotein complexes were expressed in *Escherichia coli* JM109(DE3) by co-transformation with three mutually compatible expression plasmids: pCDFDuet-1 encoding Cas11 with an N-terminal hexahistidine tag and an HRV 3C protease recognition site, pRSFDuet-1 encoding Cas5, Cas6, Cas7, Cas8 and Cas11, and pACYCDuet-1 encoding the target-specific crRNA. Cells were cultured in 2×YT medium at 37 °C with shaking until the optical density at 600 nm reached 0.6–0.8. Protein expression was induced with 0.4 mM isopropyl β-D-1-thiogalactopyranoside (IPTG), followed by incubation at 26 °C for 16 h.

Cells were harvested and lysed, and the crRNA-loaded Cascade complexes were purified by nickel-nitrilotriacetic acid (Ni-NTA) affinity chromatography. The N-terminal hexahistidine tag was removed by HRV 3C protease digestion, and the complexes were further purified by size-exclusion chromatography in buffer containing 350 mM NaCl, 1 mM dithiothreitol and 20 mM HEPES-Na (pH 7.0). The purity and apparent molecular composition of the purified Cascade complexes were evaluated by SDS–PAGE.

### RT–LAMP primer design and screening

Candidate RT–LAMP primer sets were designed within the selected target regions using PrimerExplorer V5 (Eiken Chemical Co., Ltd., Japan). PrimerExplorer V5 was used to generate LAMP primer combinations from the input target sequences, including the corresponding loop primers where applicable. Candidate primer sets were subsequently evaluated experimentally using in vitro-transcribed (IVT) RNA as the template.

Primer development was performed through sequential rounds of experimental screening. Candidate sets were first subjected to an initial screen, after which primer sets meeting the predefined selection criteria were advanced to refined screening. Selected candidates were subsequently subjected to further optimization and final kinetic evaluation.

Each primer set was evaluated in three replicate reactions containing IVT RNA (+RNA) and three matched no-template control (NTC) reactions. NTC reactions contained the complete RT–LAMP primer mixture and all other reaction components, but IVT RNA was replaced with distilled water. Primer sets were advanced only when all three of the following criteria were satisfied: (i) none of the three NTC reactions crossed the predefined fluorescence threshold (0/3 positive; Gate 1); (ii) all three +RNA reactions crossed the threshold (3/3 positive; Gate 2); and (iii) the mean time to threshold (Tt) of the three +RNA reactions was ≤15 min (Gate 3). Accordingly, amplification in even one of the three NTC reactions resulted in exclusion of that primer set from subsequent screening.

When the number of primer sets satisfying all three criteria exceeded that required for the subsequent screening round, qualifying candidates were ranked in ascending order of mean Tt. This criterion was used solely as an operational rule for candidate prioritization and was not interpreted as a statistical significance test. An additional RT–LAMP blank containing neither IVT RNA nor RT–LAMP primers was included to assess background fluorescence arising from the reaction mixture itself. The omitted RNA and primer volumes were replaced with distilled water. The RT–LAMP blank was used as a background control and was not used for the primer-specific Gate 1 selection criterion.

### Reverse transcription loop-mediated isothermal amplification and fluorescence analysis

Target RNAs were amplified by reverse transcription loop-mediated isothermal amplification (RT–LAMP) using HeatAct LAMP Master for Fluorescence (NIPPON GENE, cat. no. 311-09821). Each 10-µl reaction contained 5 µl of 2× HeatAct LAMP Master Mix, 0.08 units of AMV reverse transcriptase (NIPPON GENE, cat. no. 311-07501), 1 µl of 10× RT–LAMP primer mix, 1 µl of RNA template and nuclease-free water. The 10× primer mix contained 16 µM each of the forward and backward inner primers (FIP and BIP), 2 µM each of the forward and backward outer primers (F3 and B3), 8 µM each of the forward and backward loop primers (LF and LB), and 10 mM Tris–HCl (pH 8.0; NIPPON GENE, cat. no. 312-90061). The final primer concentrations in the reaction were therefore 1.6 µM each for FIP and BIP, 0.2 µM each for F3 and B3, and 0.8 µM each for LF and LB. Reactions were incubated at 65 °C for 40 min, with fluorescence monitored in real time using a CFX Connect Real-Time PCR Detection System (Bio-Rad). Each experiment included a no-template control (NTC), in which the RNA template was replaced with an equal volume of nuclease-free water. Unless otherwise indicated, completed RT–LAMP reaction mixtures were used directly in subsequent CONAN reactions without purification. RT–LAMP primer sequences are provided in Supplementary Table 5.

Fluorescence data were exported from the CFX Connect system and analysed using Microsoft Excel and GraphPad Prism 11 (GraphPad Software). For each reaction, baseline fluorescence was calculated as the mean fluorescence measured during the first three acquisition points. The fluorescence threshold was defined separately for each reaction as its baseline fluorescence plus 20,000 relative fluorescence units (RFU). Time to threshold (Tt) was defined as the first recorded time point at which the fluorescence exceeded this threshold. During RT–LAMP primer screening, the mean Tt for each primer set was calculated from the three RNA-positive (+RNA) replicate reactions only when all three replicates reached the threshold.

### Fluorescence-based CONAN assay

Fluorescence-based Cas3-operated nucleic acid detection (CONAN) was performed with modifications to previously described methods [17,20]. Synthetic double-stranded DNA (dsDNA) targets were prepared by annealing pairs of complementary 60-nucleotide DNA oligonucleotides synthesized by Eurofins Genomics. Equimolar amounts of the complementary oligonucleotides were mixed, heated at 95 °C for 5 min and cooled to 25 °C at a rate of 0.1 °C s⁻¹. The resulting 60-bp dsDNA targets were diluted with distilled water (NIPPON GENE, 316-90101) to working concentrations of 1 or 0.1 µM. Oligonucleotide sequences are provided in Supplementary Table 6.

All CONAN master mixes were prepared on ice. Each 10-µl reaction contained 100 nM target-specific Cascade–crRNA complex, 400 nM EcoCas3, 1 mM ATP (Takara Bio, 4041), 1 µM single-stranded DNA fluorescent reporter (5′-FAM-ATATAT-BHQ1-3′; Fasmac) and 1× CONAN buffer comprising 60 mM KCl (Nacalai Tesque, 13092-85), 10 mM MgCl₂ (Nacalai Tesque, 95812-85), 10 µM CoCl₂ (Nacalai Tesque, 08781-24) and 5 mM HEPES–KOH, pH 7.5 (Nacalai Tesque, 15639-84). For assays using synthetic DNA, 1 µl of annealed dsDNA was added to 9 µl of CONAN master mix to yield a final target concentration of 100 or 10 nM. An equal volume of nuclease-free water was used for the no-target control.

For assays coupled to RT–LAMP, 1 µl of the completed RT–LAMP product was added to 9 µl of CONAN master mix. All selected RT–LAMP primer sequences are provided in Supplementary Table 7. The corresponding no-template RT–LAMP product was used as the negative control, and the same volume of RT–LAMP product was added to all reactions within each experiment. Reactions were incubated at 37 °C in a CFX Connect Real-Time PCR Detection System (Bio-Rad), and fluorescence was measured in the FAM channel every 30 s for 20 min and reported as relative fluorescence units (RFU).

### Analysis of fluorescence-based CONAN reactions

Fluorescence data were exported from the CFX Connect Real-Time PCR Detection System and analysed using GraphPad Prism 11 (GraphPad Software). RFU values at individual time points or at the specified assay endpoint were used for comparisons. Each experiment included the corresponding no-target or no-template control. Background-corrected fluorescence was calculated by subtracting the RFU value of the corresponding no-target or no-template control from that of each sample.

### Lateral-flow-based CONAN assay

Lateral-flow-based CONAN was performed in a total reaction volume of 20 µL. Each reaction contained 50 nM target-specific Cascade–crRNA complex, 200 nM EcoCas3, 1 mM ATP, 250 nM single-stranded DNA lateral-flow reporter (5′-FITC-ATATAT-biotin-3′; Fasmac) and 1× CONAN buffer, as described above. A 17-µl CONAN master mix was combined with 2 µl of completed RT–LAMP product and 1 µl of the lateral-flow reporter. Reactions were incubated at 37 °C for 10 min. Following the CONAN reaction, 80 µL of assay buffer supplied with the Milenia GenLine HybriDetect kit (Milenia Biotec, Giessen, Germany) was added to each reaction. A lateral-flow strip was inserted into the reaction tube, and the result was recorded after 3 min.

### Analytical specificity and cross-reactivity testing

Analytical specificity was evaluated using synthetic in vitro-transcribed (IVT) RNAs at 1 × 10^5^ copies per RT–LAMP reaction. For ANDV, SNV, LANV and CHOV RNAs were amplified using the ANDV-specific RT–LAMP primer set and analysed using the ANDV-specific CONAN assay without blinding. Cross-reactivity among the filovirus assays was evaluated using BDBV, EBOV, SUDV, TAFV and MARV IVT RNAs. RT–LAMP reactions were prepared by an experimenter who retained the sample identities, whereas the experimenter performing the subsequent CONAN assays was blinded to sample identity and well position.

An RT–LAMP blank containing neither IVT RNA nor RT–LAMP primers was included as a negative background control. CONAN reactions with a mean normalized positivity score (P̄) ≥ 0.5 were classified as positive and those with P̄ < 0.5 as negative. Cross-reactivity was defined as a positive classification for a non-cognate viral RNA. T1/2 was analysed separately as a kinetic metric and was not used interchangeably with the P̄-based classification criterion.

### Lyophilization and storage of CONAN reagents

CONAN reagents were lyophilized with modifications to a previously described procedure [17]. The Cas3–Cascade protein components and CONAN reaction-buffer components were lyophilized separately. For the protein formulation, EcoCas3 and target-specific crRNA-loaded ecCascade were mixed in a stabilizing solution containing 250 mM trehalose, 20 mM HEPES–NaOH (pH 7.0) and 200 mM NaCl. Each 2-µL aliquot contained Cas3 at 0.45 µg µL⁻¹ and crRNA-loaded ecCascade at 0.40 µg µL⁻¹. The aliquots were pre-frozen at −80 °C and subsequently lyophilized using a freeze-drying system equipped with a dry chamber (Tokyo Rikakikai Co., Ltd., Tokyo, Japan).

CONAN reaction-buffer components were prepared and lyophilized separately. After reconstitution, the buffer contained 5 mM HEPES–KOH (pH 7.0), 60 mM KCl, 10 mM MgCl₂ and 10 µM CoCl₂ and combined with ATP. Lyophilized protein and buffer components were reconstituted with distilled water immediately before use, reporter and target nucleic acid to obtain the indicated final reaction conditions.

For storage-stability testing, lyophilized Cas3–Cascade preparations were stored at 25 °C and evaluated after 11, 30 and 70 days using an EGFP-targeting crRNA-loaded Cascade complex. Lyophilized CONAN reaction-buffer preparations were evaluated in parallel for up to 57 days. Freshly prepared Cas3–Cascade and reaction-buffer preparations were used as comparators. Collateral-cleavage activity was measured at 37 °C using an EGFP-containing plasmid target and a FAM–BHQ1 single-stranded DNA reporter.

### Operation of the portable CONAN-SWIFT system

CONAN-SWIFT was operated as a portable two-step workflow combining lyophilized RT-LAMP amplification, lyophilized Cas3-based CONAN detection and lateral-flow assay (LFA) readout. Lyophilized RT-LAMP reagents (HeatAct LAMP Master; NIPPON GENE) were reconstituted with the sample and incubated at 65 °C for 30 min using the battery-operated CAMP isothermal amplification device (NIPPON GENE). Following RT-LAMP amplification, an aliquot of the amplification product was transferred to the reconstituted lyophilized CONAN reaction containing EcoCas3 and the target-specific crRNA-loaded ecCascade complex, prepared as described above, and incubated at 37 °C for 5 min in the CAMP device. LFA readout was performed using the same procedure described above for the lateral-flow-based CONAN assay. A result was considered valid when the control line was visible, and target detection was indicated by the appearance of a test line. The workflow was performed without a cold chain, centrifugation or mains power, and was additionally evaluated at ambient temperatures of 24 °C and 37 °C. The complete step-by-step procedure, including reagent reconstitution, reaction volumes, incubation conditions, transfer steps and visual interpretation criteria, is provided in Supplementary Protocol 3 and demonstrated in Supplementary Video 1.

### Preparation and handling of EBOV ΔVP30

EBOV ΔVP30 was generated as previously described [23]. Briefly, VeroVP30 cells were maintained in Eagle’s minimum essential medium (MEM) supplemented with 10% fetal calf serum (FCS), L-glutamine, vitamins, non-essential amino acids and 5 μg ml⁻¹ puromycin (Sigma-Aldrich, St. Louis, MO, USA). EBOV ΔVP30 was propagated in VeroVP30 cells in MEM supplemented with 2% FCS. Virus-containing material was harvested 6 days after infection at a multiplicity of infection of 0.1 and partially purified by ultracentrifugation at 27,000 rpm for 2 h through a 20% sucrose cushion. The resulting viral pellet was resuspended in sterile phosphate-buffered saline (PBS) and stored at −80 °C until use. Viral titres were determined by a focus-forming assay in confluent VeroVP30 cells, with infected foci visualized using an antibody against the viral VP40 protein. The prepared EBOV ΔVP30 stock was diluted and spiked into human whole blood or wastewater samples before analysis by CONAN-SWIFT.

### Whole-blood samples

Human whole blood obtained from healthy donors was used to evaluate the performance of CONAN-SWIFT in blood samples. Whole blood was diluted fourfold with nuclease-free water before analysis. For analytical sensitivity experiments, synthetic in vitro-transcribed ANDV, BDBV and EBOV RNAs were added at the indicated input copy numbers and analysed by RT-LAMP followed by CONAN detection and lateral-flow readout. EBOV ΔVP30 was spiked into whole-blood samples obtained independently from three healthy donors and analysed using the same workflow. Unspiked blood from each donor was processed in parallel as a negative control. Human RPP20 was detected in parallel as an endogenous internal control to confirm successful sample processing and amplification. Lateral-flow results were interpreted visually as described above.

### Wastewater sample collection and processing

Wastewater samples were collected from a wastewater treatment plant and sewer system of an international airport in Japan. Wastewater samples were supplemented with MgCl₂ to a final concentration of 25 mM, filtered through an electronegative mixed cellulose ester membrane (Millipore; 0.8-µm pore size) following the previously described protocol [33]. The membrane was divided into four pieces, three of which were subjected to direct nucleic acid extraction using the RNeasy PowerWater Kit (QIAGEN, Hilden, Germany), with a Precellys Evolution homogenizer for the bead-beating step. DNase treatment was omitted to allow recovery of both RNA and DNA. The nucleic acid extracts were subsequently subjected to RT-LAMP–CONAN analysis.

### Detection of viral RNA from wastewater-derived nucleic acid extracts using RT-LAMP–CONAN

For analytical sensitivity experiments, synthetic BDBV and EBOV RNAs were added at 10¹–10⁵ copies per reaction to nucleic acid extracts from concentrated municipal wastewater (MW1) or airport wastewater (AW1), or to nuclease-free water as an RNA-only control, and analysed by RT-LAMP followed by CONAN detection and lateral-flow readout. Values were reported as the number of positive reactions among three independent reactions at each input level (Extended Data Fig. 7b).

For experiments with biologically contained Ebola virus, EBOV ΔVP30 was spiked into airport and urban wastewater samples after the concentration step. The spiked samples were then subjected to CONAN-SWIFT analysis. Pepper mild mottle virus (PMMoV), a highly abundant wastewater-associated virus [34], was detected in parallel as a per-sample process control. Lateral-flow results were interpreted visually as described above.

### Use of artificial-intelligence-assisted language tools

During preparation of this manuscript, the authors used ChatGPT (OpenAI) for English-language editing and structural revision. The authors reviewed and edited all generated text and take full responsibility for the content of the manuscript.

### Statistical analysis

Data were analysed using GraphPad Prism 11 (GraphPad Software). For comparisons involving three or more groups, one-way analysis of variance followed by Dunnett’s multiple-comparisons test was used for data meeting the assumptions of parametric testing. The Kruskal–Wallis test followed by Dunn’s multiple-comparisons test was used for non-parametric data. Comparisons between two groups were performed using a two-tailed Student’s t-test or Mann–Whitney U-test, depending on the distribution of the data. Statistical significance was defined as P < 0.05. The statistical test, number of independent replicates, definition of the center and error bars, and exact P values are provided in the corresponding figure legends.

## Acknowledgements

This research was primarily supported by the Japan Agency for Medical Research and Development (AMED) under Grant No. JP223fa627001 (Y.K). T.M. was also supported by AMED under Grant Nos. JP24fk0108690, JP223fa727002, JP223fa627006, JP23bm1223009 and JP26bk0104207; JSPS KAKENHI Grant No. JP23H00367; and the JST Program on Open Innovation Platform for Industry–Academia Co-creation (COI-NEXT), Grant No. JPMJPF2010. S.T. was supported by AMED under Grant Number JP223fa627001 (UTOPIA Next-Generation Diagnostic Tool Discovery Program) and the Environment Research and Technology Development Fund (grant number JPMEERF20255RB1 and JPMEERF20265001) of the Environmental Restoration and Conservation Agency provided by Ministry of the Environment of Japan. M.K. was supported by AMED under Grant No. 26fk0108713h0003. K.Y. was supported by AMED under Grant No. JP23ck0106807, JP26ck0106096 and JSPS KAKENHI Grant No. 24K02010. The authors thank Dr. Made Sandhyana Angga for his assistance with wastewater sample preparation, Miss. Hiromi Taniguchi for her experimental assistance, and Mr. Tsubasa Obo for his taking video protocol.

## Author contributions

T.M. and Y.K. conceived and supervised the study and wrote the manuscript. J.N. and K. Miyazaki performed the principal experiments, analyzed the data and compiled the results. S.T. and M.K. designed and conducted the experiments using wastewater. K. Mikamo, L.M., H.A., R.H., S.I. and K.Y. assisted with the CONAN-SWIFT experiments. T.K. and J.I. developed the CONAN-SWIFT Designer. K.T., S.K. and Y.M. expressed and purified the Cascade proteins. M.I. and P.J.H. prepared and purified the replication-deficient Ebola virus. All authors reviewed and approved the manuscript.

## Data availability

All data supporting the findings of this study are available within the Article and its Supplementary Information. Additional data are available from the corresponding author upon reasonable request. Sequence information for the synthetic RNA templates, crRNAs, synthetic DNA targets and RT-LAMP primers used in this study is provided in Supplementary Tables 1–7. The viral genome sequences used for crRNA target selection were retrieved from publicly available records in the Pathoplexus SeqSet (https://pathoplexus.org/) and the NCBI Nucleotide database. The accession numbers, sequence identifiers and source databases for all sequences used in this analysis are provided in Supplementary Table 8.

## Code availability

The source code for CONAN-SWIFT Designer is available at [https://github.com/kimihira-tetsutaro-code/conan-swift-designer.git]. All other custom code used in this study is available from the corresponding author upon reasonable request.

## Supplementary Data legends

**Supplementary Protocol 1 | CONAN-SWIFT Designer user guide.**

User guide for sequence-guided design and prioritization of CONAN-SWIFT candidates, including FASTA input, candidate scoring and filtering, optional NCBI BLAST off-target analysis, and result export.

**Supplementary Protocol 2 | Standard RT-LAMP–CONAN protocol.**

Detailed laboratory protocol for two-step RT-LAMP amplification followed by Cas3-operated nucleic acid detection (CONAN), including fluorescence- and lateral-flow-based readouts, reaction compositions, transfer steps, controls and interpretation criteria.

**Supplementary Protocol 3 | Portable CONAN-SWIFT workflow.**

Step-by-step protocol for portable CONAN-SWIFT testing using lyophilized RT-LAMP and CONAN reagents, the battery-operated CAMP isothermal device, and lateral-flow visual readout. The workflow includes reagent reconstitution, 30-min RT-LAMP amplification, 5-min CONAN detection and 3-min lateral-flow development, enabling visual detection within 40 min.

**Supplementary Protocol 4 | Extraction-free CONAN-SWIFT analysis of human whole blood.**

Protocol for direct CONAN-SWIFT analysis of human whole blood without nucleic acid extraction, including endogenous RPP20 and spiked viral targets.

**Supplementary Protocol 5 | CONAN-SWIFT analysis of wastewater samples.**

Protocol for CONAN-SWIFT analysis of wastewater, including sample concentration, nucleic acid extraction and detection of endogenous and spiked viral targets.

**Supplementary Video 1 | Portable operation of the CONAN-SWIFT system.**

Representative video demonstrating the complete portable CONAN-SWIFT workflow using lyophilized reagents and the battery-operated CAMP isothermal device. Lyophilized RT-LAMP reagents were reconstituted and incubated for 30 min, followed by transfer of the amplified product to the reconstituted lyophilized CONAN reaction and incubation for 5 min. The reaction was then analyzed using a lateral-flow assay, with visual interpretation after 3 min. The complete workflow was completed within 40 min. The corresponding step-by-step procedure is provided in Supplementary Protocol 3.

---

**Extended Data Fig. 1.**
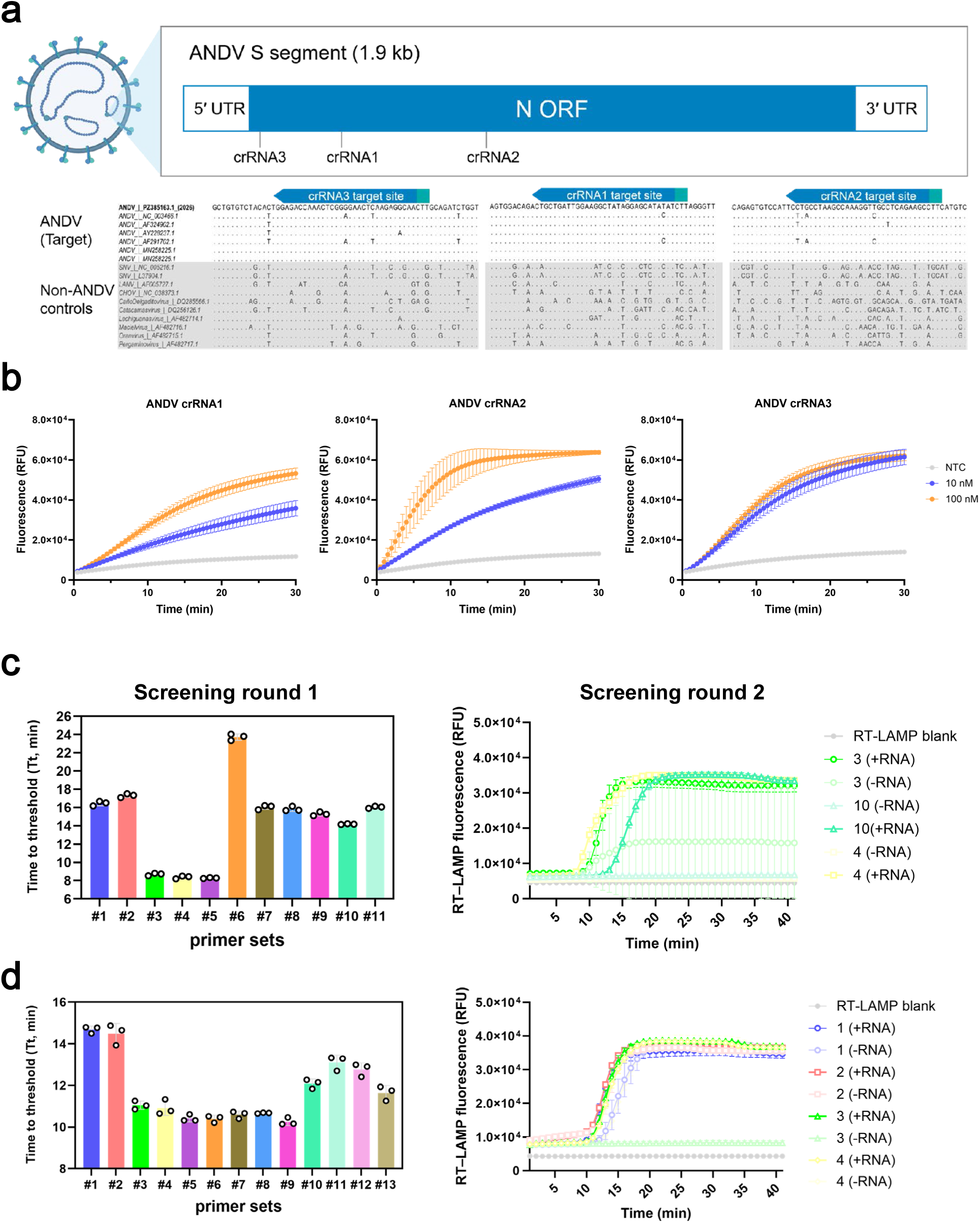
Sequence selection and experimental screening of candidate components for the ANDV RT-LAMP–CONAN assay. **a,** Multiple-sequence alignments of the three candidate crRNA target regions in ANDV and related hantaviruses. Dots indicate nucleotide identity to the ANDV reference sequence; crRNA spacer regions and AAG protospacer-adjacent motifs (PAMs) are indicated. **b,** Fluorescence-based CONAN screening of three candidate ANDV crRNAs using synthetic double-stranded DNA targets at the indicated concentrations. c,d, RT-LAMP primer screening for regions encompassing the ANDV crRNA2 (c) and crRNA3 (d) target sites. The dashed line indicates the 15-min time-to-threshold (Tt) screening criterion. Data in b are mean ± s.d. of three replicate reactions. NTC, no-target control; RFU, relative fluorescence units.

**Extended Data Fig. 2.**
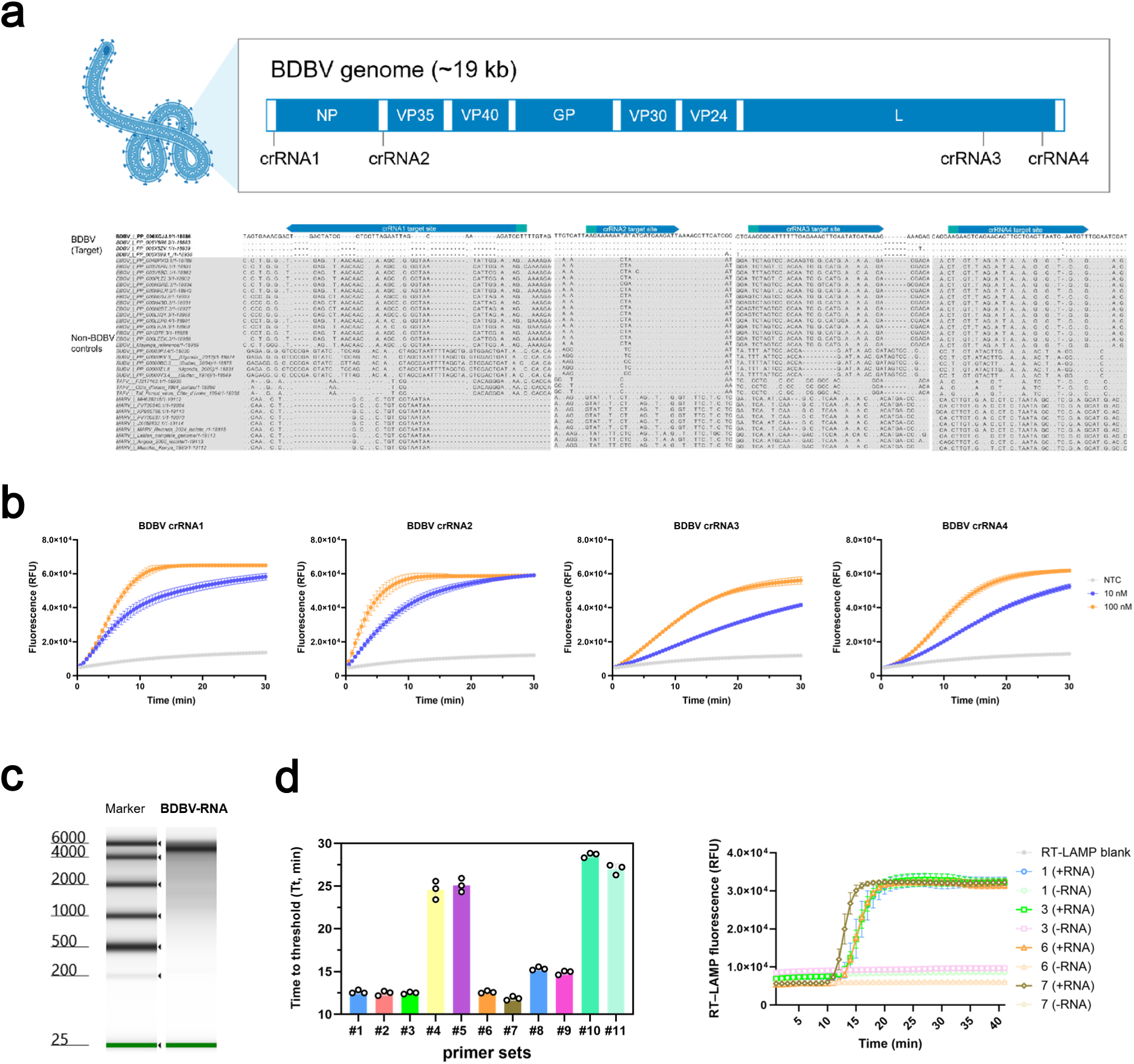
Sequence selection and experimental screening of candidate components for the BDBV RT-LAMP–CONAN assay. **a,** Schematic representation of the Bundibugyo virus (BDBV) genome and the candidate target region used for assay development, and multiple-sequence alignment of genomic regions encompassing four candidate BDBV crRNA target sites across BDBV and other medically important filoviruses. Dots indicate nucleotide identity to the BDBV reference sequence; crRNA spacer regions and protospacer-adjacent motifs (PAMs) are indicated. **b,** Fluorescence-based CONAN screening of candidate BDBV crRNAs using synthetic double-stranded DNA targets at the indicated concentrations, resulting in selection of crRNA2 for subsequent assay development. Data are shown as mean ± s.d. of three replicate reactions. NTC, no-target control; IVT, in vitro transcription; RFU, relative fluorescence units. **c,** Representative TapeStation analysis of purified synthetic in vitro-transcribed (IVT) BDBV RNA used for assay development. RNA marker sizes are indicated in nucleotides (nt). **d,** RT-LAMP primer screening for regions encompassing the BDBV target sites. The dashed line indicates the 15-min time-to-threshold (Tt) screening criterion. Data in d are mean ± s.d. of three replicate reactions. NTC, no-target control; RFU, relative fluorescence units.

**Extended Data Fig. 3.**
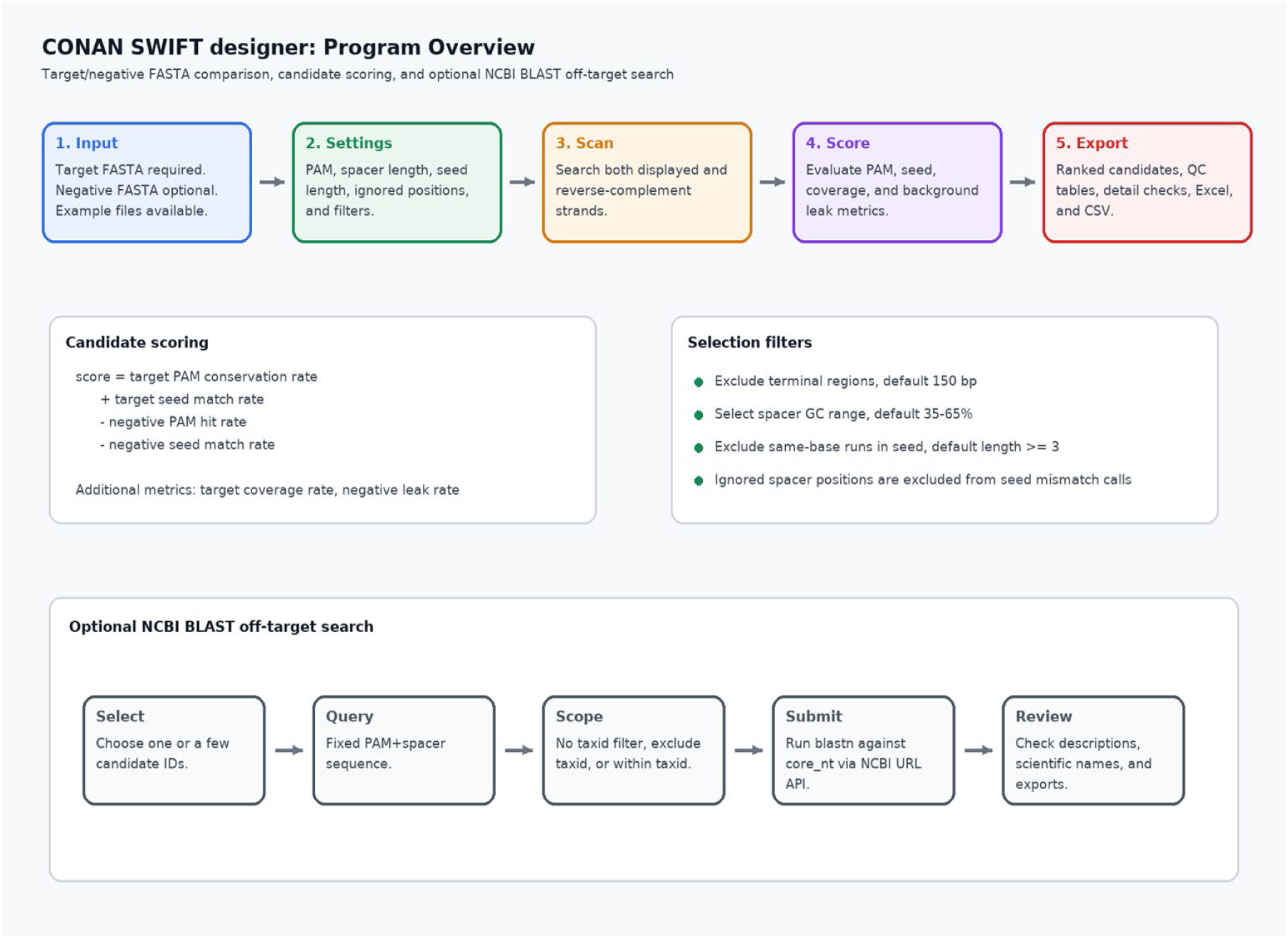
Overview of the CONAN-SWIFT Designer workflow. Schematic overview of the CONAN-SWIFT Designer workflow for sequence-guided crRNA selection. Target and optional negative FASTA sequences are used to identify and rank candidate crRNAs according to PAM conservation, target coverage and sequence specificity, with configurable selection filters and optional NCBI BLAST off-target analysis. Ranked candidates and quality-control information can be exported for downstream assay development.

**Extended Data Fig. 4.**
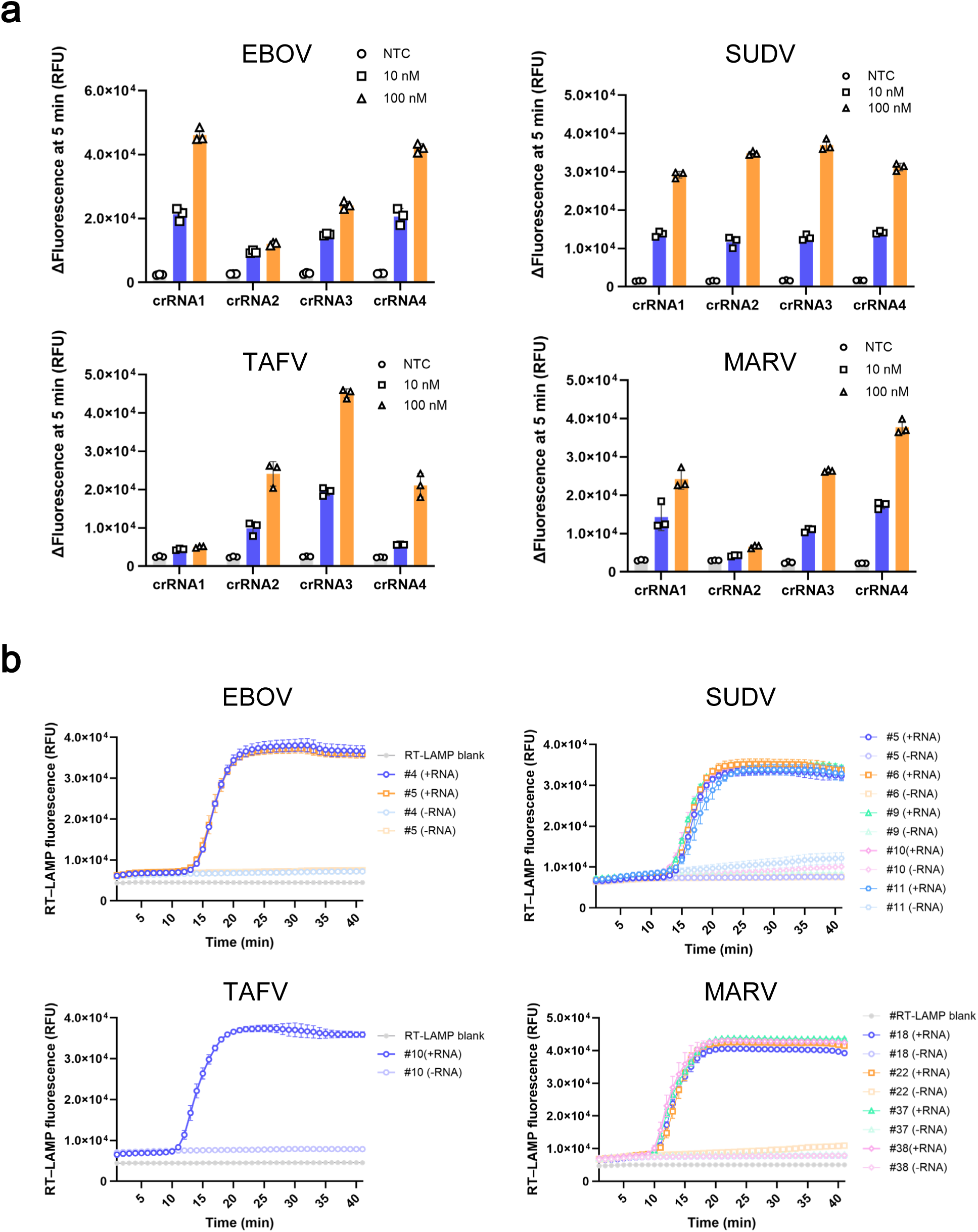
Optimization and extension of CONAN-SWIFT assays to multiple filoviruses. **a,** Fluorescence-based screening of candidate crRNAs for Ebola virus (EBOV), Sudan virus (SUDV), Taï Forest virus (TAFV) and Marburg virus (MARV) using synthetic double-stranded DNA targets. **b,** RT-LAMP primer screening for EBOV, SUDV, TAFV and MARV using the corresponding synthetic in vitro-transcribed (IVT) viral RNAs. Candidate primer sets were evaluated on the basis of amplification kinetics and specificity against no-RNA controls. Data are shown as individual replicate reactions with summary statistics as indicated. NTC, no-target control; IVT, in vitro transcription.

**Extended Data Fig. 5.**
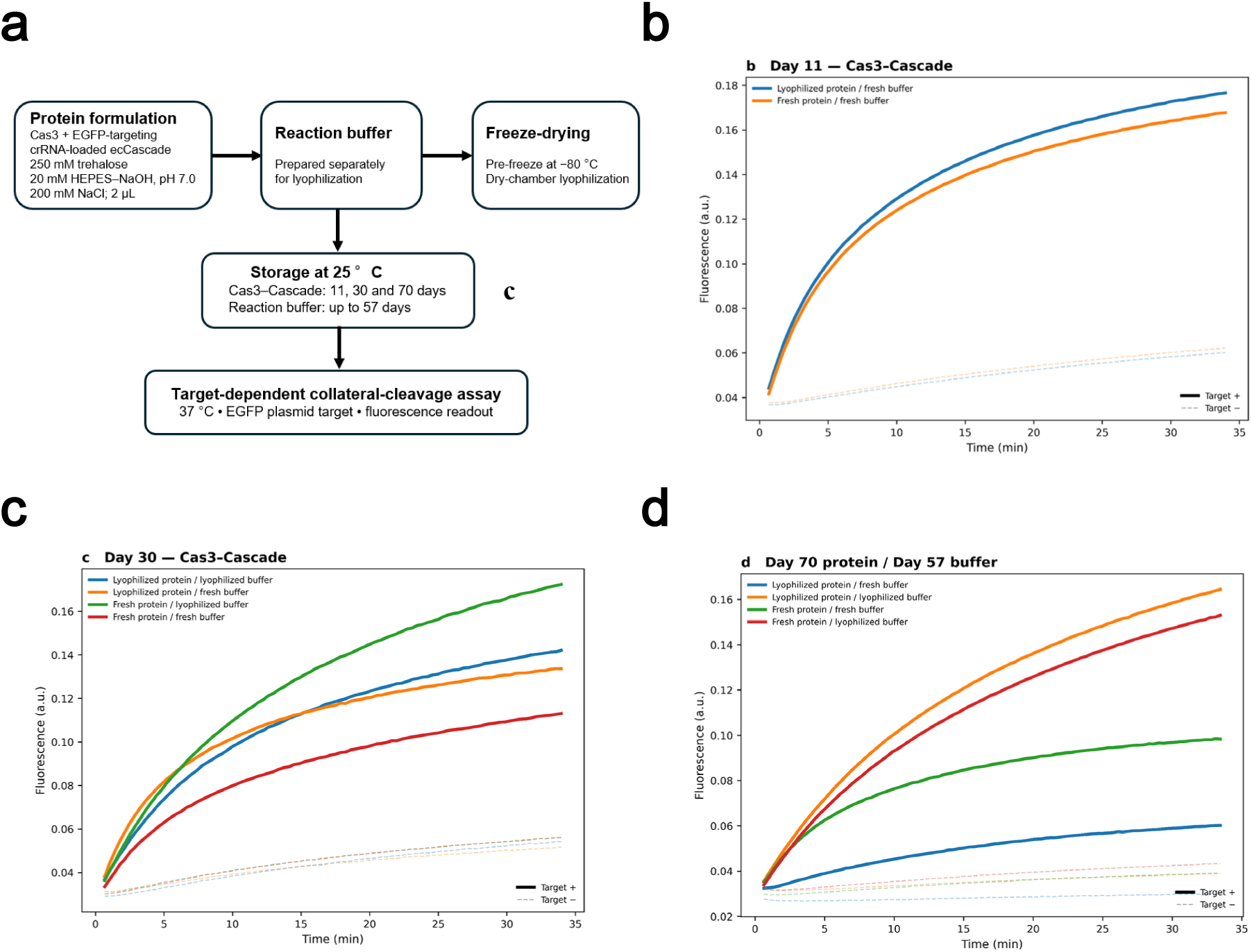
Ambient-temperature stability of lyophilized CONAN reagents. **a,** Schematic of the lyophilization and storage-stability workflow. Cas3 and EGFP-targeting crRNA-loaded ecCascade and CONAN reaction buffer were lyophilized separately and stored at 25 °C before activity testing. **b,** Comparison of freshly prepared and lyophilized Cas3–Cascade after 11 days of storage. **c,** Comparison of fresh and lyophilized Cas3–Cascade and reaction-buffer components after 30 days of Cas3–Cascade storage. **d,** Corresponding comparison after 70 days of Cas3–Cascade storage; the lyophilized reaction buffer had been stored for 57 days. Target-positive reactions are shown as solid lines and no-target controls as dashed lines. Lyophilized Cas3–Cascade retained target-dependent collateral-cleavage activity after storage at 25 °C for up to 70 days.

**Extended Data Fig. 6.**
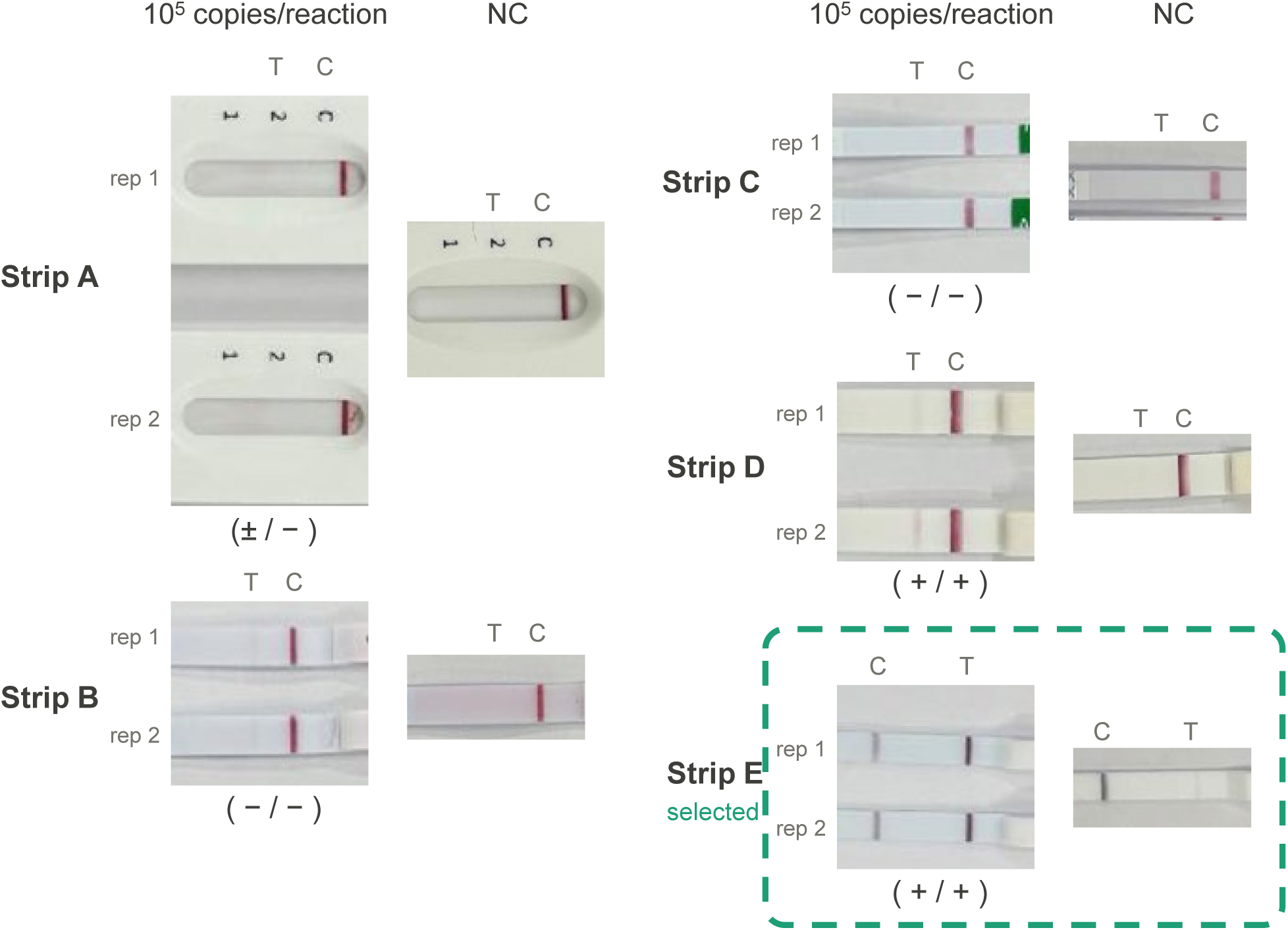
Comparison of lateral-flow strips for visual detection of CONAN reaction products. Five commercially available lateral-flow strips were evaluated using CONAN reaction products generated from 10⁵ copies of RPP20 RNA and corresponding no-template controls (NTCs). Two independent reactions were tested for each condition. Visual scores indicate clearly visible (+), faint or ambiguous (±), or absent (−) test lines. The strip selected for subsequent experiments is outlined. T, test line; C, control line; NTC, no-template control.

**Extended Data Fig. 7.**
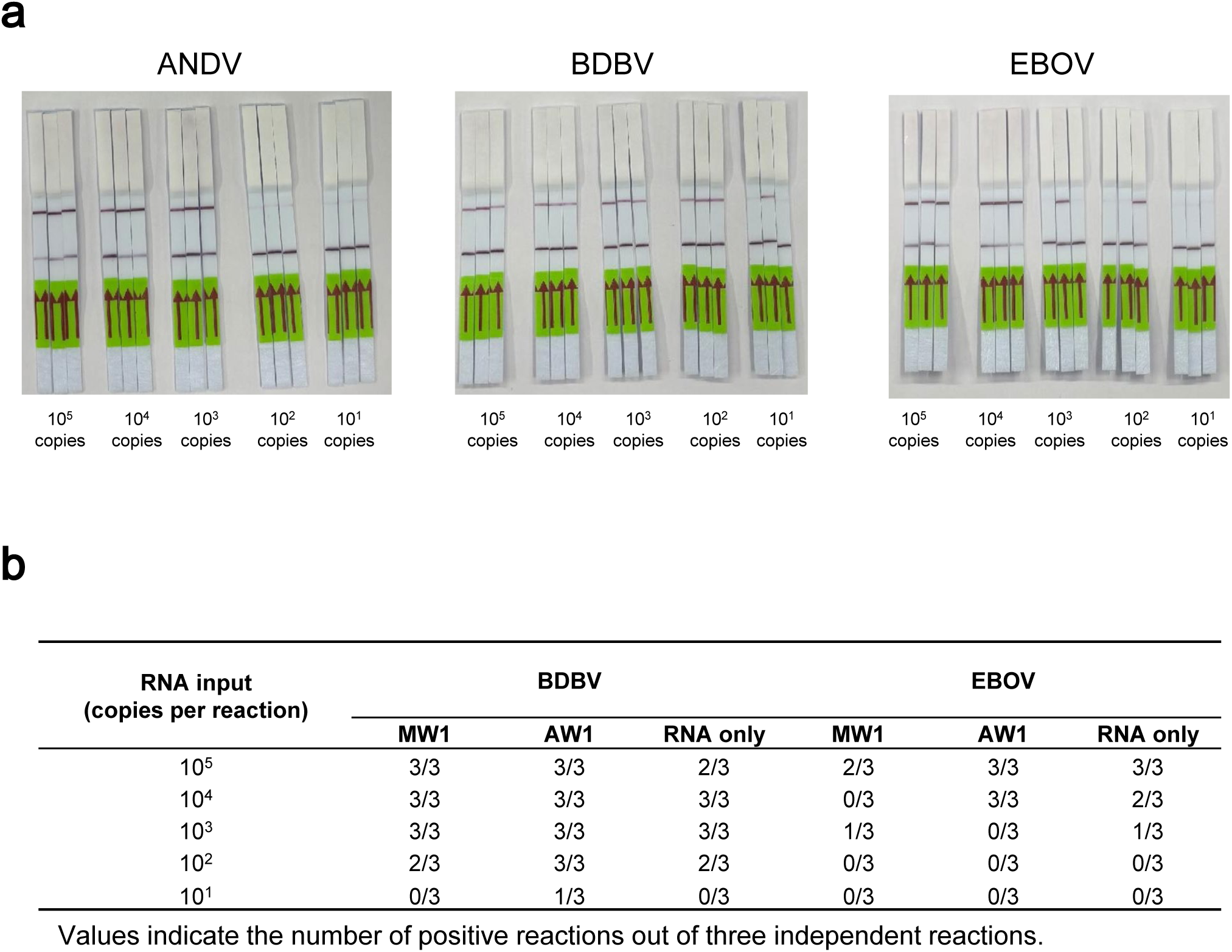
Analytical sensitivity of CONAN-SWIFT in human whole blood and wastewater matrices. **a,** Representative lateral-flow readouts from analytical sensitivity experiments using synthetic in vitro-transcribed (IVT) ANDV, BDBV and EBOV RNAs spiked into human whole blood at the indicated input copy numbers. Samples were processed by RT-LAMP followed by CONAN detection and lateral-flow readout. Strips are arranged in order of decreasing RNA input. A visible test line indicates target detection. **b,** Detection frequency of synthetic BDBV and EBOV RNAs in wastewater-derived nucleic acid extracts, as determined by fluorescence-based RT-LAMP-CONAN. Viral RNA was added at 10¹–10⁵ copies per reaction to RNA extracted from concentrated municipal wastewater (MW1) or airport wastewater (AW1), or to nuclease-free water (RNA only), and analysed by RT-LAMP–CONAN. Values indicate the number of positive reactions among three independent reactions at each input level. C, control line; T, test line; NTC, no-template control.

